# *Cbfb2* gene dosage programs the differential lymphoid lineage developmental potential of fetal and adult hematopoietic progenitors

**DOI:** 10.64898/2026.03.02.708762

**Authors:** Alyssa Berthelette, Kaelie Newell, Quan Phan, Ichiro Taniuchi, Joonsoo Kang, Michela Frascoli

**Author notes:** Correspondence (J.K.), (M.F.). These authors contributed equally (Joonsoo Kang, Michela Frascoli).

## Abstract

Innate-like lymphocyte subsets are generated predominantly during early-life windows, yet the mechanisms that restrict their development in adulthood remain unclear. Here we identify *Cbfb2* gene dosage as a quantitative regulator of stage-specific lymphoid potential. We show that reduction of CBFβ2 levels unlocks fetal-like competence in adult hematopoietic progenitors, enabling robust generation of IL-17–producing γδ T (Tγδ17) cells. Although *Cbfb2* haploinsufficiency minimally alters steady-state transcription, chromatin profiling of H3K4me3 revealed promoter-level changes in adult lymphoid-primed multipotent progenitors consistent with altered developmental priming. In adult bone marrow chimeras, *Cbfb*^+*/2m*^ progenitors efficiently generated functional Vγ2^+^ Tγδ17 cells in lymph nodes and skin, and restoring *Cbfb2* expression suppressed this capacity, establishing a dosage-dependent mechanism. Using an optimized in utero transplantation system, we further demonstrate that fetal niches amplify this latent competence and selectively favor IL-17–committed γδ T cell differentiation over conventional αβ T cell output. *Notch1* haploinsufficiency enhanced Tγδ17 generation and phenocopied the effect of CBFβ2 dosage reduction, linking quantitative NOTCH1 signaling to innate-like lymphocyte developmental programming. Together, these findings reveal that fetal versus adult lymphopoiesis is governed by quantitative tuning of RUNX:CBFβ activity and uncover unexpected plasticity in adult hematopoiesis controlled by transcription factor dosage.

## Introduction

Although numerous immune cell subsets are produced throughout an individual’s lifespan, certain specialized lymphocytes are only generated within a restricted window of time. Immune cells that exclusively or primarily originate early in life include innate and innate-like lymphocytes (diverse γδ and αβ T cell subsets, B-1 B cells, and innate lymphoid cells) and they play dual function at mucocutaneous barriers: they promote tissue homeostasis and repair, especially in early life, and are the first responders to tissue infections. Adult bone marrow (BM) hematopoietic stem cells (HSCs) exhibit limited capacity to produce most types of innate-like lymphocytes, even after tissue insults, and as a result, older animals tend to have diminished numbers of innate-like lymphocytes that can lead to exaggerated tissue inflammatory responses^1–3^. Although it is well-established that distinct gene programs govern fetal and adult HSCs^4–6^, and transcriptional regulators that can confer fetal lymphoid lineage developmental potential in adult progenitors have been described^7,8^, it is unknown to what degree adult HSCs and multi-potent progenitors are fixed from generating innate lymphocytes of fetal origin.

Dermal lymphocytes expressing the γδ TCR and secreting IL-17 (Tγδ17) serve as a model for innate-like T cell development and function. Tγδ17 cells are produced in the late fetal and early neonatal stages and are crucial for maintaining skin barrier homeostasis^9^. Dermal Tγδ17 cells originate from embryonic hematopoietic tissues^10,11^ and are long-lived in the skin, not relying on constant reinforcements from the thymus. Adult BM progenitors do not efficiently contribute to the pool of skin Tγδ17 cell^10,12–15^. The molecular mechanisms that govern the differential ability of fetal and adult hematopoietic progenitors to generate Tγδ17 cells and other innate-like lymphocyte subsets have not been fully elucidated. A simplified model posits that either the tissue niches program age-dependent progenitor developmental potential in trans or that early life-defined progenitors have cell intrinsic differentiation programs that are distinct from adult progenitors. While the search for age-specific factors have naturally focused on distinct gene signatures of fetal versus adult hematopoiesis^16–18^, ectopic expression of such factors alone fails to reprogram adult progenitors to generate fetal-origin innate-like lymphocytes^19^. An overlooked possibility is the role of core transcriptional regulator gene dosage in imposing age-dependent developmental potential of hematopoiesis throughout life. The RUNX (Runt-related transcription factor) family of transcription factors (TFs) is a central regulator of hematopoiesis^20^, including cell lineage fate choice. The Core Binding Factor subunit β (CBFβ), an evolutionarily conserved protein that forms a heterodimeric complex with RUNX proteins, is the obligatory partner of RUNX TFs^21^. CBFβ stabilizes the DNA binding domain of RUNX and enhances its DNA binding affinity, resulting in increased chromatin and transcription regulatory activity. By controlling the expression of cell lineage-specific genes, RUNX:CBFβ complex plays a crucial role in instituting definitive embryonic hematopoiesis^18,19^ from the endothelium^22,23^. The mammalian *Cbfb* gene can produce two major protein isoforms by alternative splicing, CBFβ1 and CBFβ2, which differ in their C-terminal sequences^24^. While CBFβ1 isoform has been shown to have only a minor, non-redundant function in lymphopoiesis^24,25^, CBFβ2 confers homing capacity to prethymic progenitors during embryogenesis^24^ and it is absolutely required for the generation of all T cell subsets. While there has been some intriguing data suggesting that the quantity of CBFβ as a factor in distinct lymphoid lineage differentiation in adults^26^ the finding has not coalesced into an understanding of gene dosage as a principal parameter of context-dependent hematopoietic system construction. Consistent with this concept, recent work has shown that RUNX factors act as quantitative and context-dependent regulators of early lymphoid fate, with dynamic shifts in RUNX binding, RUNX cofactor recruitment, and RUNX protein availability controlling the premature or delayed activation of T cell-identity and innate lymphoid cell programs^27^. These findings highlight RUNX dosage and functional partner availability as critical parameters shaping lineage specification. However, whether quantitative differences in RUNX:CBFβ complex activity could have broad consequences for early-life restricted immune subset differentiation remains undetermined.

CBFβ transcriptional regulation is poorly understood. In zebrafish, *Cbfb* expression is regulated by NOTCH1^28^, and binding sites for the NOTCH effector RBP-Jκ are present within *Cbfb* regulatory regions in bone marrow multipotent progenitors^29^. More recently, a comprehensive study demonstrated that NOTCH signaling drives a functional conversion of RUNX transcription factors to initiate the T-lineage program^30^. NOTCH signaling represents a fundamental regulatory pathway that shapes T cell development and lymphoid lineage decisions. The seminal discovery that NOTCH1 activity is essential for T cell specification^31^ established the broader principle that evolutionarily conserved morphogenetic pathways, including NOTCH, WNT, Hedgehog, and BMP/TGFβ, govern hematopoietic fate decisions throughout life^32^. Early studies demonstrated that *Notch1* heterozygosity skews αβ versus γδ lineage choice toward γδ T cells, suggesting that quantitative modulation of NOTCH activity influences early T lineage bifurcation^31^. Although NOTCH signaling is indispensable for the emergence of definitive HSCs^33,34^ in the embryonic Aorta-Gonad-Mesonephros region and is reacquired at the LMPP/ETP stages^35^, the specific features of NOTCH1 activity that endow fetal progenitors with a unique competence to generate innate-like lymphocytes, including Tγδ17 cells, remain poorly understood.

In this study, we found that *Cbfb2* haploinsufficiency renders adult HSCs and progenitors permissive for the development of Tγδ17 and other innate-like lymphocytes. In adult LMPPs, reduced amounts of RUNX:CBFβ2 complex impose fetal stage epigenome states without overt changes in the transcriptome. In utero transplantation studies further revealed that this progenitor cell-intrinsic alteration in CBFβ2 dosage, and consequent changes in developmental potential, is amplified in fetal developmental niches. Importantly, Tγδ17 cells generated from adult *Cbfb2* heterozygous BM are functionally competent, as they elicited dominant IL-17 responses in both imiquimod (IMQ)-induced psoriasis-like dermatitis and fungal skin colonization with *Malassezia furfur*. Together, these findings identify *Cbfb2* gene dosage as a principal determinant of fetal versus adult lymphopoiesis that underpins dermal immune homeostasis. Consistent with this model, *Notch1* heterozygosity similarly unlocks fetal-like lymphoid potential in adult progenitors.

## Results

### CBFβ2 haploinsufficiency limits precursor supply without impairing Tγδ17 cell maturation

CBFβ2 is required for the differentiation of early thymic progenitors (ETPs) and their homing to the thymus^24^. Absence of this isoform impairs T cell development in general, with a block in γδ T cell development^24^. In order to assess the effect of *Cbfb2* heterozygosity on the development of late fetal/early neonatal IL-17 secreting γδ T cells (Tγδ17), we analyzed mice with a targeted mutation in the splice donor signal of *Cbfb2*, causing 50% reduction of CBFβ2 isoform in heterozygous mice (*Cbfb*^*+/2m*^) and complete loss in homozygous mice (*Cbfb*^*2m/2m*^), in comparison to littermate wild-type mice (*Cbfb*^*+/+*^)^24^. Dermal Tγδ17 cells arise from CD4^-^CD8^-^ double negative (DN), stage 1 (CD44^+^CD25^-^), subset d (DN1d) thymic precursors^36^. While DN1d cells arise at E16.5, expression of the TF *Sox13*, that underpins generation of neonatal Vγ2^+^CCR6^+^Scart2^+^ Tγδ17 cells^11^ (Garman nomenclature; Vγ2=Vγ4 in Tonegawa designation^37^), peaks at day 7 after birth. In heterozygous mice, overall T cell development was not grossly altered with most DN thymic precursor subsets in the normal range (**Fig. S1xx**). However, DN1d cells, while present in *Cbfb2*^*+/2m*^ fetuses (from E16.5 to 0 day old), were reduced in numbers in neonatal (7 day old) *Cbfb2*^*+/2m*^ mice (**Fig. 1A, B; Supp. Fig. 1A, B**). In *Cbfb*^*2m/2m*^ mice early thymic progenitors (DN1a/b) were present at higher frequencies, although reduced in absolute numbers, but DN1d cells were depleted at all ages (**Fig. 1A, B; Supp. Fig. 1A, B**). Unexpectedly, despite the paucity of DN1d cells after birth, *Cbfb*^*+/2m*^ mice supported normal numbers of Vγ2^+^ cells throughout the late embryonic and early neonatal developmental stages (**Fig. 1C**), and frequencies of mature Vγ2^+^ Tγδ17 thymocytes were comparable to wild-type mice (**Fig. 1D**). Skin draining lymph nodes (sLN) and dermis were also populated with similar numbers of mature Vγ2^+^ Tγδ17 cells in adult heterozygous mice (**Fig. 1E, F**). This result suggests that in postnatal *Cbfb2*^*+/2m*^ mice, other sources of Vγ2^+^ Tγδ17 cells besides DN1d precursors exist.

**Figure 1.**
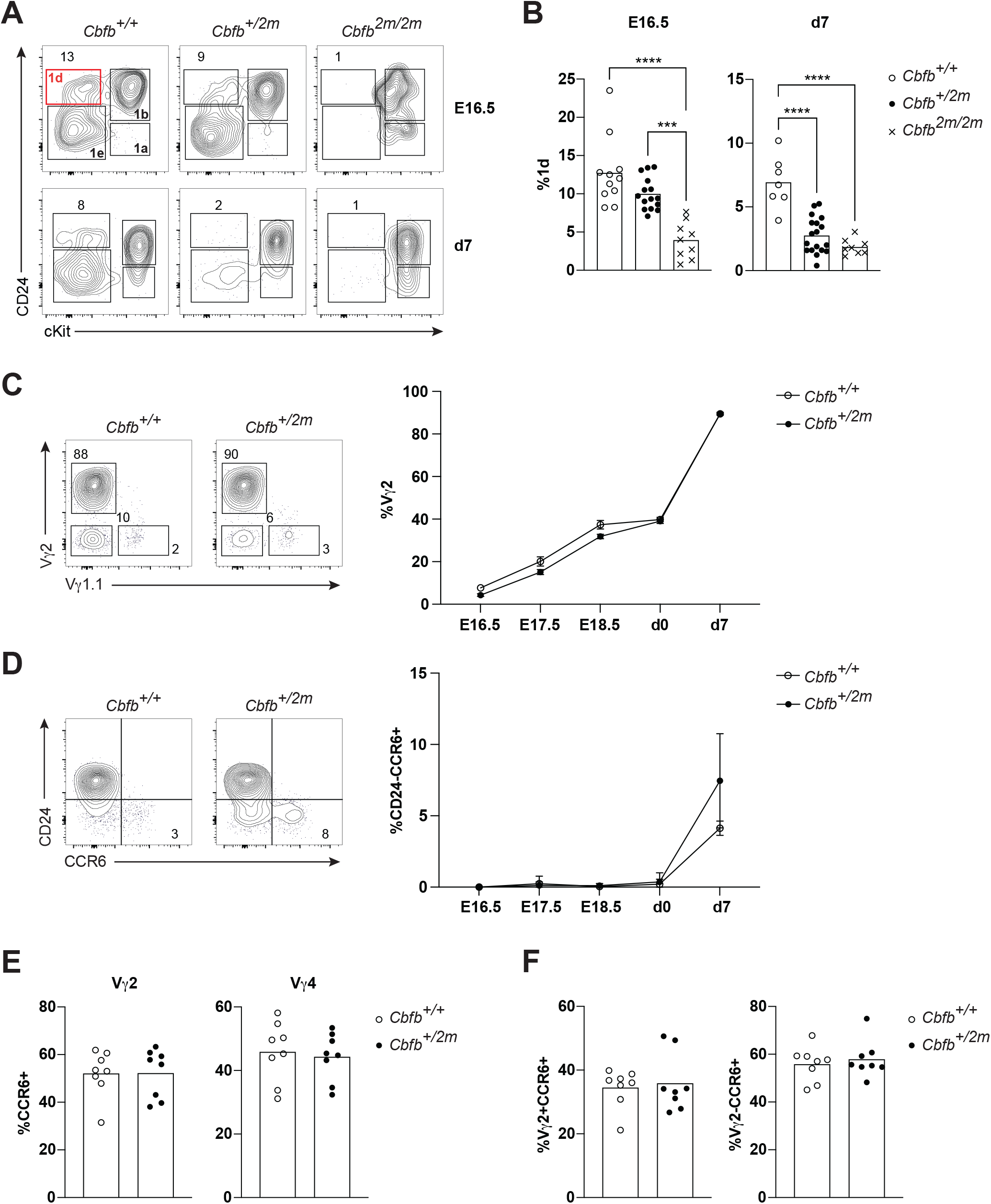
*Cbfb2* gene dosage controls neonatal Tγδ17 cell differentiation in the thymus. (A) Representative flow cytometric contour plots depicting expression of CD24 and cKit in DN1 (CD44^+^CD25^-^ DN) thymocytes from *Cbfb*^*+/+*^, *Cbfb*^*+/2m*^, and *Cbfb*^*2m/2m*^ mice analyzed at E16.5 and d7 post-partum. (B) Summary of the compiled flow cytometric data of A (n?3 mice per genotype). P values determined by two-way ANOVA. (C) Representative flow cytometric profiles of γδT cell subsets in the thymus of 7 days old *Cbfb*^*+/+*^and *Cbfb*^*+/2m*^ neonate mice (left). Summary of frequencies of Vγ2 thymocytes at the indicated time points (n?4 mice per group) (right). (D) Representative flow cytometric profiles of CD24 and CCR6 expression on Vγ2^+^ cells in the thymus of 7 days old *Cbfb*^*+/+*^and *Cbfb*^*+/2m*^ mice (left). Summary of frequencies of mature Vγ2^+^ Tγδ17 thymocytes (CD24^-^CCR6^+^) at the indicated time points (n?4 mice per group) (right). (E) Summary of frequencies of CCR6 expression on Vγ2^+^ and Vγ1.1^-^Vγ2^-^ cells in skin-draining LN (sLN) of adult *Cbfb*^*+/+*^and *Cbfb*^*+/2m*^ mice (n=8 mice per genotype). (F) Summary of frequencies of Vγ2^+^CCR6^+^ and Vγ2^-^CCR6^+^ cells in the dermis of adult *Cbfb*^*+/+*^and *Cbfb*^*+/2m*^ mice (n=8 mice per genotype). Data in (A-F) are representative of at least 3 independent experiments. Each symbol represents one mouse. *P* values determined by one-way ANOVA (B), two-way ANOVA (C-D) or unpaired t-test (E-F).

### Bone marrow progenitors from CBFβ2 haploinsufficient mice are capable of reconstituting functional Vγ2^+^ Tγδ17 cells

Tγδ17 cells are poorly reconstituted by adult BM in transplantation models^10,12–15^. A significant generation of Tγδ17 cells was only observed in adult host when neonatal thymocytes were injected together with BM^13^, an indication that the reconstitution potential might reside in the perinatal thymus when the Vγ2^+^ Tγδ17 physiologically arises. To test the possibility that another source of Vγ2^+^ Tγδ17 cells in *Cbfb*^*+/2m*^ mice distinct from DN1d precursors exists, we assessed whether CBFβ2 haploinsufficiency rendered a permissive state for adult BM progenitors to differentiate into Tγδ17 cells. BM chimeras were generated using either *Cbfb*^*+/+*^ and *Cbfb*^*+/2m*^ BM (CD45.2) cells to reconstitute irradiated C57BL/6 (CD45.1) mice. As expected, *Cbfb*^*+/+*^ BM cells minimally contributed to the Vγ2^+^ Tγδ17 pool with the majority made up of partially radio-resistant host Tγδ17 cells; in contrast heterozygous *Cbfb*^*+/2m*^ (HET) BM cells efficiently generated Vγ2^+^ Tγδ17 cells in skin draining lymph nodes (sLN) (**Fig. 2A, B; Supp. Fig. 2A**) and dermis (**Fig. 2C, D; Supp. Fig. 2A**). Fetal-origin Vγ4^+^ Tγδ17 cells with distinct developmental gene circuits were minimally, and comparably, reconstituted by either donor types (**Supp. Fig. 2B-E**), suggesting that CBFβ2 haploinsufficiency specifically impacts Vγ2^+^ Tγδ17 cells. sLN and skin Vγ2^+^ Tγδ17 cells derived from *Cbfb*^*+/2m*^ BM cells produced IL-17 when restimulated (**Fig. 2E**), showing their intact functionality.

**Figure 2.**
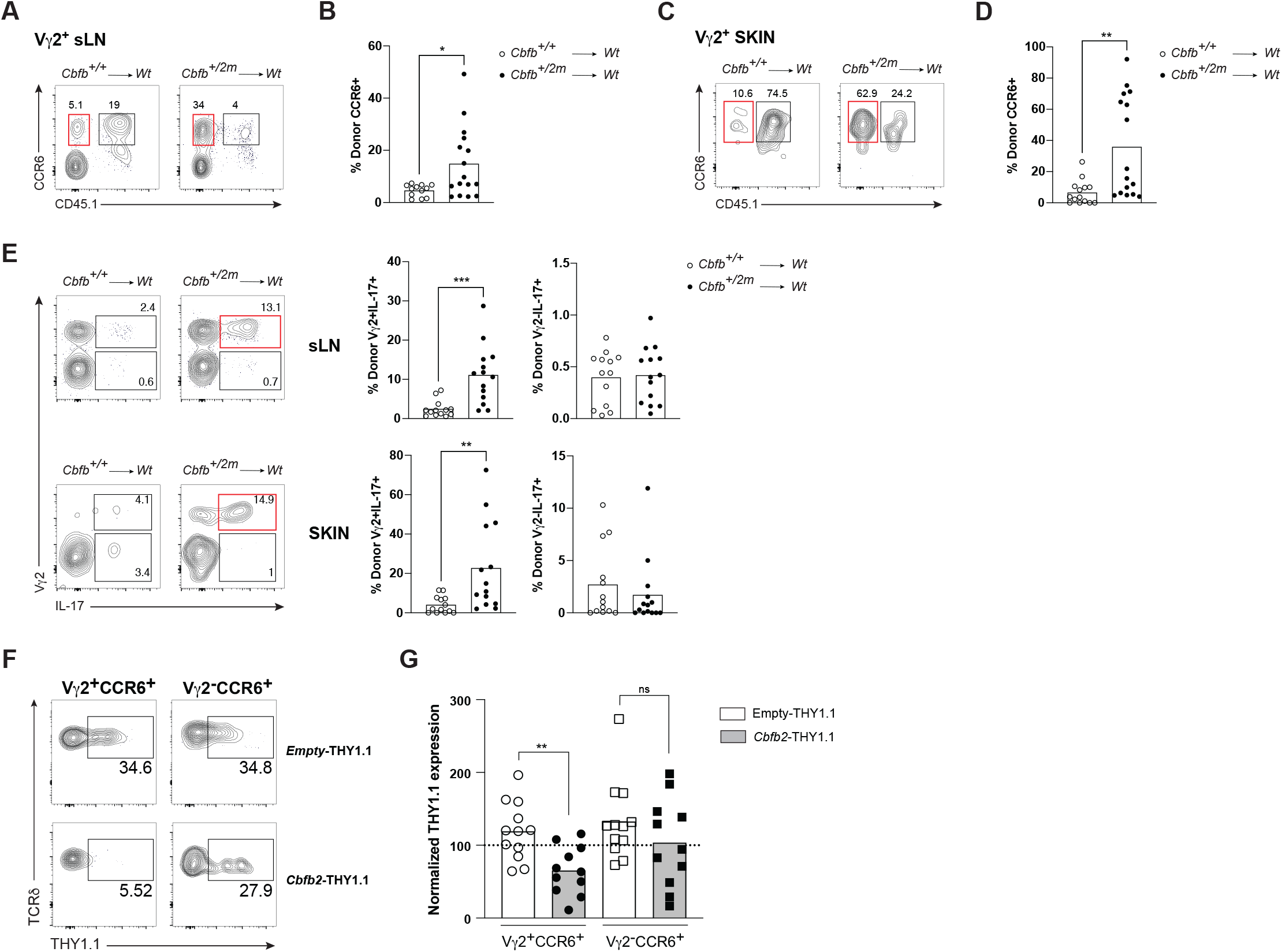
Efficient Tγδ17 cell reconstitution with *Cbfb2* haploinsufficient BM cells. (A) Representative flow plots of CCR6 expression among Vγ2^+^ cells in BM chimera sLN. (B) Compiled frequencies of CCR6^+^ donor cells among Vγ2^+^ cells from (A) (n?12 mice per group). (C) Representative flow plots of CCR6 expression among Vγ2^+^ cells in BM chimera skin dermis. (D) Compiled frequencies of CCR6^+^ donor cells among Vγ2^+^ cells from (C) (n?12 mice per group). (E) Representative intracellular staining for IL-17 amongst γδT cells in BM chimera sLN and skin dermis (left) and summary of the compiled frequencies of donor Vγ2^+^IL17^+^ and donor Vγ2^-^IL17^+^ cells (right). (n?13 mice per group). (F) Representative profiles of transduced cells (indicated as THY1.1) among Vγ2^+^CCR6^+^ and Vγ2^-^CCR6^+^ cell subsets in cervical LNs of animals reconstituted with either empty vector (top panels) or *Cbfb2* virus transduced BM progenitors (bottom panels). (G) Summary of Vγ2^+^CCR6^+^ and Vγ2^-^CCR6^+^ cell subsets after normalization of THY1.1 expression (n?11 mice per group). Red highlight gates indicate Vγ2^+^CCR6^+^ (A-C) and Vγ2^+^IL17^+^ (E) cells of donor origin (CD45.1^-^). Data in (A-G) are representative of at least 3 independent experiments. Each symbol represents one mouse. *P* values determined by unpaired t-test.

To address the formal possibility that the adult *Cbfb*^*+/2m*^ BM cells had accumulated alterations other than CBFβ2 proteins, we tested whether increased amounts of CBFβ2 alone in heterozygous BM progenitors will impede Tγδ17 cell generation. We transduced adult *Cbfb*^*+/2m*^ HSCs with retroviruses carrying *Cbfb2* and generated radiation BM chimeras. Overexpression of *Cbfb2* in heterozygous BM cells resulted in the suppression of Vγ2^+^ Tγδ17 cell generation in LNs of reconstituted mice when compared to the control HET BM transduced with an empty vector (**Fig. 2F, G**). In the αβ T cell compartment, the conventional CD4 and CD8 αβ T cell subsets were indistinguishable upon *Cbfb2* overexpression in HET progenitors. The IL-17+ Mucosa-Associated Invariant T (MAIT17) subsets originating from *Cbfb2* retrovirus transduced HET HSCs showed a trend to lower production, but this was not statistically significant, with high variability (**Supp. Fig. 2F**). Together, these results indicate that a reduction in the amounts of CBFβ2 confer neonatal-type Tγδ17 cell generative potential to adult BM progenitors.

To further evaluate the effector capacity of Vγ2^+^ Tγδ17 cells generated from HET BM, we assessed their responses in two well-established inflammatory models in which Vγ2^+^ Tγδ17 cells are known primary drivers: psoriasis-like dermatitis^14,38,39^ and fungal skin colonization^40^. Psoriasis was induced by topical application with Imiquimod (IMQ), a TLR7/8 agonist, for five consecutive days. As expected, this treatment resulted in increased ear thickness indicative of overt inflammatory responses (**Fig. 3A**). Flow-cytometry analysis of lymphocytes isolated from sLN and dermis and treated with Brefeldin A alone to capture cytokine secretion without ex-vivo re-stimulation, revealed increased IL-17 production from donor-derived Vγ2^+^ Tγδ17 cells in both sLN (**Fig. 3B**) and skin dermis (**Fig. 3C**) compared to untreated controls. A similar increase in IL-17 secretion was also observed in donor-derived Vγ4^+^ Tγδ17 cells in both sLN (**Supp. Fig. 3A**) and skin dermis (**Supp. Fig. 3B**). For the fungal skin colonization model, chimeric mice were epicutaneously associated with *Malassezia furfur* for seven consecutive days (**Fig. 3D**). In this setting, donor-derived Vγ2^+^ Tγδ17 cells from both sLN and skin dermis showed increased IL-17 secretion compared to Olive Oil-treated controls (**Fig. 3E, F**). No differences were observed for donor-derived Vγ4^+^ Tγδ17 cells (**Supp. Fig. 3C, D**). These results show that Vγ2^+^ Tγδ17 cells derived from adult HET BM progenitors are indistinguishable in function from WT counterparts.

**Figure 3.**
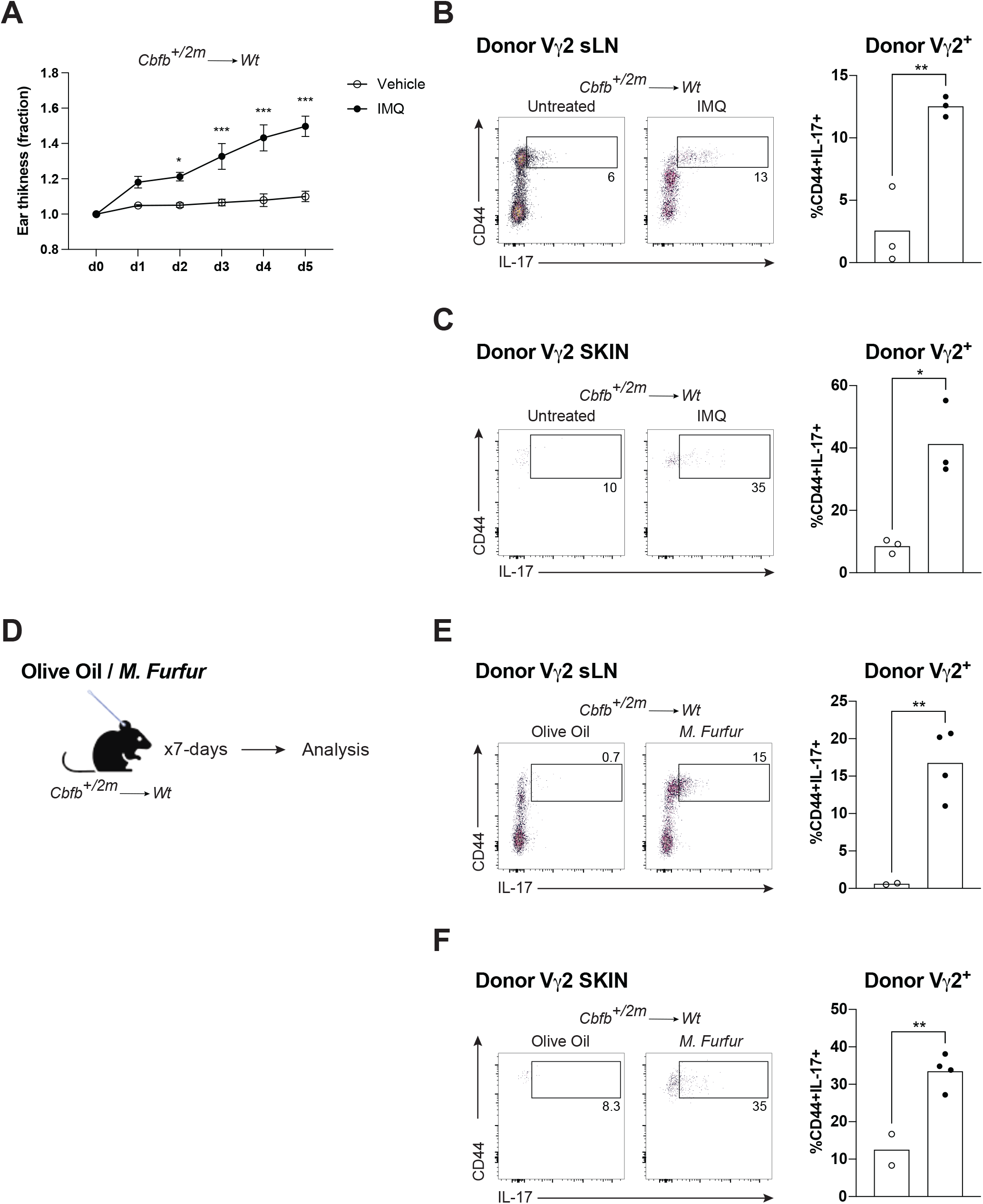
CBFβ2 haploinsufficiency enables adult bone marrow derived Vγ2^+^ Tγδ17 cells to mount IL-17–driven skin inflammatory responses. (A) Relative ear skin thickness (set at 1 at day 0) measurements for 6 days from untreated or IMQ-treated chimeric mice (n=3 mice per group). (B) Representative flow plots of IL-17 secretion among donor Vγ2^+^ cells in BM chimera sLN after IMQ treatment and corresponding compiled frequencies (n=3 mice per group). (C) Representative flow plots of IL-17 secretion among donor Vγ2^+^ cells in BM chimera skin dermis after IMQ treatment and corresponding compiled frequencies (n=3 mice per group). (D) Schematic of fungal skin colonization model with *M. Furfur*. (E) Representative flow plots of IL-17 secretion among donor Vγ2^+^ cells in BM chimera sLN after fungal colonization and corresponding compiled frequencies (n?2 mice per group). (F) Representative flow plots of IL-17 secretion among donor Vγ2^+^ cells in BM chimera skin dermis after fungal colonization and corresponding compiled frequencies (n?2 mice per group). Each symbol represents one mouse. *P* values determined by two-way ANOVA (A) or by unpaired t-test (B, C, E, F).

### *Cbfb2* gene dosage minimally alters adult progenitor transcriptomes while remodeling chromatin states

The classical model of HSC differentiation in the FL and BM, entails serial stages of differentiation to generate lymphocytes (MultiPotent Progenitors (MPPs), Lymphoid-Primed MPPs (LMPPs), Common Lymphoid Progenitors (CLPs), and ETPs). To begin to identify the CBFβ2-modulated gene circuits that specify Tγδ17 cell differentiation, we analyzed the transcriptional profiles of HSCs and lymphoid progenitors (LMPPs and CLPs) isolated from adult BM of *Cbfb*^*+/+*^ and *Cbfb*^*+/2m*^ mice. We performed Ultra Low Input (ULI) RNAseq analysis of ex-vivo sorted HSCs (Lin^-^cKit^+^Sca1^+^Flt3^-^), LMPPs (Lin^-^ cKit^+^Sca1^+^Flt3^+^) and CLPs (Lin^-^cKit^int^Sca^int^Flt3^+^IL7Ra^+^). *Cbfb*^*+/+*^ and *Cbfb*^*+/2m*^ hematopoietic progenitor transcriptomes largely overlapped, and principal component analysis showed that the three populations segregated mainly by cellular subset, rather than genotype (**Fig. 4A**). LMPPs were the most different with *Cbfb2* gene dosage (**Fig. 4A, B**), and many genes implicated in embryonic hematopoiesis were increased in expression in *Cbfb*^*+/2m*^ LMPPs (**Fig. 4C**). Among these, the known *Cbfb2*-regulated *Ccr9* expression^24^ was decreased sharply (**Fig. 4D**). The expression of *Cbfb* was decreased to 50% of *Cbfb*^*+/+*^ in *Cbfb*^*+/2m*^ LMPPs while the expression of *Runx* genes was unaltered (**Fig. 4D**). The established Vγ2^+^ Tγδ17 thymic progenitor gene signature was not differentially expressed in *Cbfb*^*+/2m*^ LMPPs, indicating that *Cbfb2* haploinsufficiency does not overtly alter known lymphoid progenitors in adult BM cells at the transcriptional level (**Supp. Fig. 4**).

**Figure 4.**
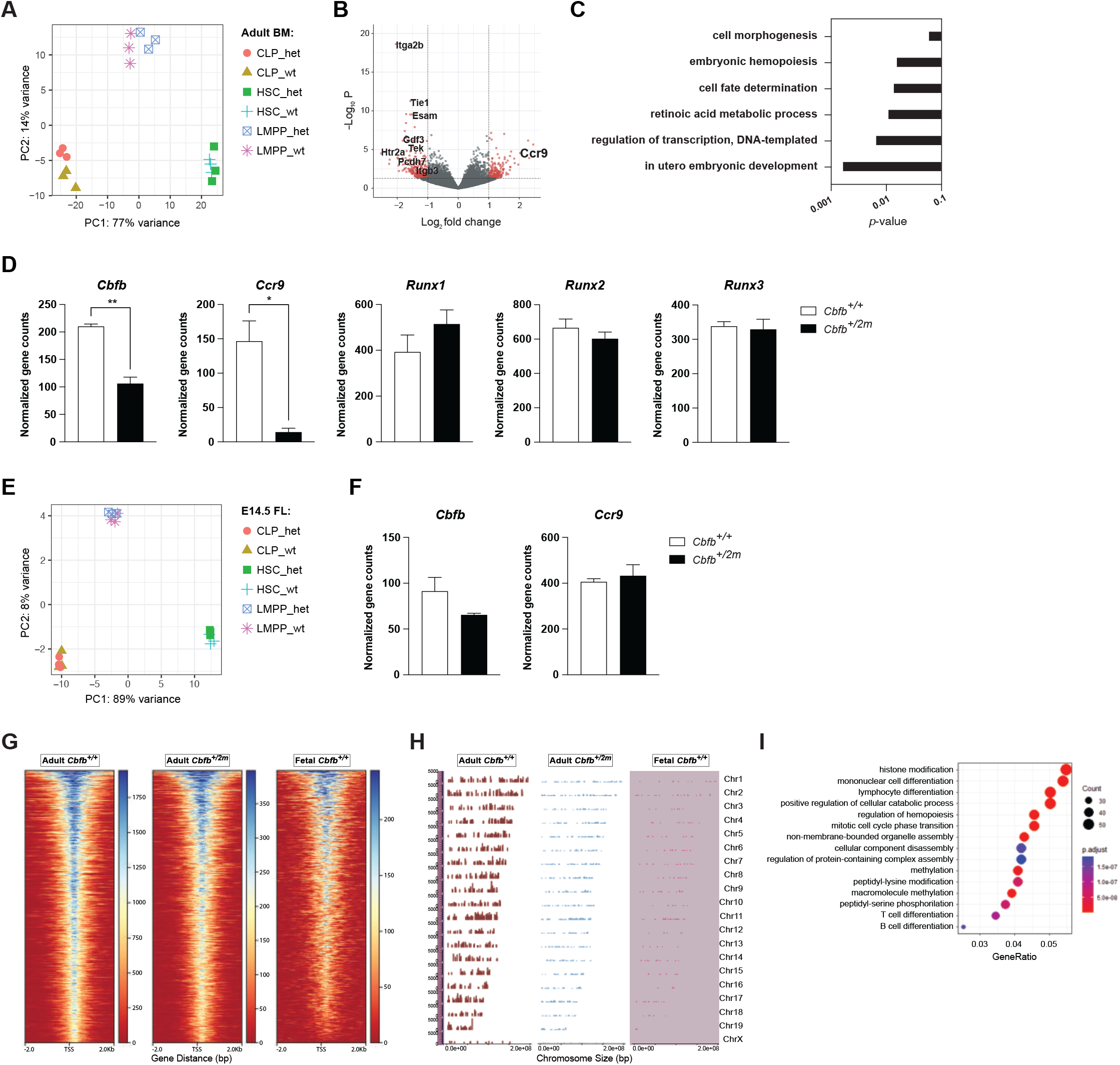
Adult *Cbfb*^*+/2m*^ LMPPs retain largely adult transcriptional profiles but acquire a fetal-like epigenetic landscape. (A) Ultra Low Input (ULI)-RNAseq Principal Component Analysis analysis of Hematopoietic Stem Cells (HSCs), Lymphoid-primed MultiPotent Progenitors (LMPPs), and Common Lymphoid Progenitors (CLPs) sorted from adult BM of *Cbfb*^*+/+*^and *Cbfb*^*+/2m*^ mice. (B) Volcano plot of differentially expressed genes of LMPPs from *Cbfb*^*+/+*^and *Cbfb*^*+/2m*^ mice, with >2-fold increase in mean expression and *P*<0.05. (C) Biological processes associated with differentially expressed genes in LMPPs from *Cbfb*^*+/2m*^ mice, based on Gene Ontology (GO) analysis with *P*<0.05. (D) Select differentially expressed genes between *Cbfb*^*+/+*^and *Cbfb*^*+/2m*^ BM-derived LMPPs, based on gene counts. (E) Ultra Low Input (ULI) RNAseq Principal Component Analysis analysis of Hematopoietic Stem Cells (HSCs), Lymphoid-primed MultiPotent Progenitors (LMPPs), and Common Lymphoid Progenitors (CLPs) sorted from E14.5 FL of *Cbfb*^*+/+*^and *Cbfb*^*+/2m*^ mice. (F) Select differentially expressed genes between *Cbfb*^*+/+*^and *Cbfb*^*+/2m*^ FL-derived LMPPs, based on gene counts. (G) Peak-centered heatmaps of CUT&RUN H3K4me3 binding profiles in LMPPs sorted from adult *Cbfb*^*+/+*^, adult *Cbfb*^*+/2m*^ and fetal *Cbfb*^*+/+*^ mice. (H) Coverage Plots of differential H3K4me3 peaks across the genomes, comparing LMPPs from adult *Cbfb*^*+/+*^, adult *Cbfb*^*+/2m*^ and fetal *Cbfb*^*+/+*^ mice. (I) GO analysis of genes associated with significantly different H3K4me3 peaks in adult *Cbfb*^*+/2m*^/*Cbfb*^*+/+*^ LMPPs. Samples were analyzed in biological triplicates for ULI-RNAseq or in duplicates for CUT&RUN. *P* values determined by the Wald test (B), modified Fisher Exact test EASEscore (C, I) or unpaired t-test (D, F).

The transcription profiles of the fetal counterpart subsets isolated from the E14.5 FL, showed an even tighter overlap between *Cbfb*^*+/+*^ and *Cbfb*^*+/2m*^ cells (**Fig. 4E**), an indication that, overall, the mutation did not have a major effect on the transcriptional landscape of fetal cell subsets. The expression of *Cbfb* was only partially reduced in *Cbfb*^*+/2m*^ LMPPs compared to wild-type mice and *Ccr9* expression was comparable. Together, these data suggest that *Cbfb2* gene dosage has a more pronounce impact on adult hematopoietic progenitors, especially LMPPs.

Considering that transcriptional differences between *Cbfb*^*+/+*^ and *Cbfb*^*+/2m*^ hematopoietic progenitor subsets are minimal, and that epigenetic remodeling events serve as the driver for the majority of cell differentiation processes, we performed a CUT&RUN assay to track the histone post-translational modification H3K4me3 in sorted FL and BM LMPPs. Globally, the patterns of the active chromatin mark of *Cbfb*^*+/2m*^ BM LMPPs were distinct from those of *Cbfb*^*+/+*^ BM LMPPs and converged toward that of FL cells (**Fig 4G, H**). Gene Ontology (GO) pathway analysis of the differentially marked active loci revealed that four of the top 15 most significantly altered categories were associated with hematopoiesis and lymphopoiesis pathways (**Fig. 4I**). These findings support a direct *Cbfb2* dose-dependent programming of lymphopoiesis at the chromatin level as the key determinant of distinct fetal versus adult hematopoiesis. Taken together, our data support the hypothesis that the generation of Tγδ17 cells is driven by a CBFβ2 dose-dependent epigenetic reprogramming of lymphoid progenitors.

### *Cbfb2* haploinsufficient adult BM progenitors effectively generate Vγ2^+^ Tγδ17 cells in utero

Currently, there are no in vivo cell transfer systems to directly assess the developmental potential of hematopoietic progenitors within a physiological fetal environment. To address this, we optimized an in utero transplantation (IUT) assay^41,42^ in which progenitors isolated from adult BM were injected into the fetal liver of E14.5 embryos without host preconditioning (**Fig. 5A**). Dams were allowed to deliver at term, and offsprings were analyzed for immune cell composition. To define the baseline performance of the IUT system, enriched Lin^−^c-Kit+ progenitors from adult WT BM were transferred into E14.5 fetal livers. In the thymus, donor-derived cells contributed at modest but reproducible levels, with comparable reconstitution of TCRδ+ and ILC (TCRβ−TCRδ−) compartments and relatively lower contribution to TCRβ+ cells (**Fig. 5B**). In contrast, in sLN donor chimerism was highly skewed: TCRβ+ and DN populations were minimally reconstituted, whereas TCRδ+ cells constituted the dominant donor-derived lineage (**Fig. 5B**). Thus, when adult BM progenitors develop in a fetal niche, the assay preferentially supports γδ T cell output over conventional αβ T cell differentiation in the periphery. Within the γδ T cell compartment of both thymus and sLN, donor-derived cells were preferentially enriched in IL-17–committed subsets compared to their IFNγ-producing counterparts (**Supp. Fig. 5A, B**). In contrast, analysis of αβ T cell populations in both thymus and sLN did not reveal preferential expansion of naïve or activated subsets (**Supp. Fig. 5C, D**). These results indicate that adult WT BM progenitors are not inherently blocked from generating early life Tγδ17 cells if allowed to differentiate in the fetal environment.

**Figure 5.**
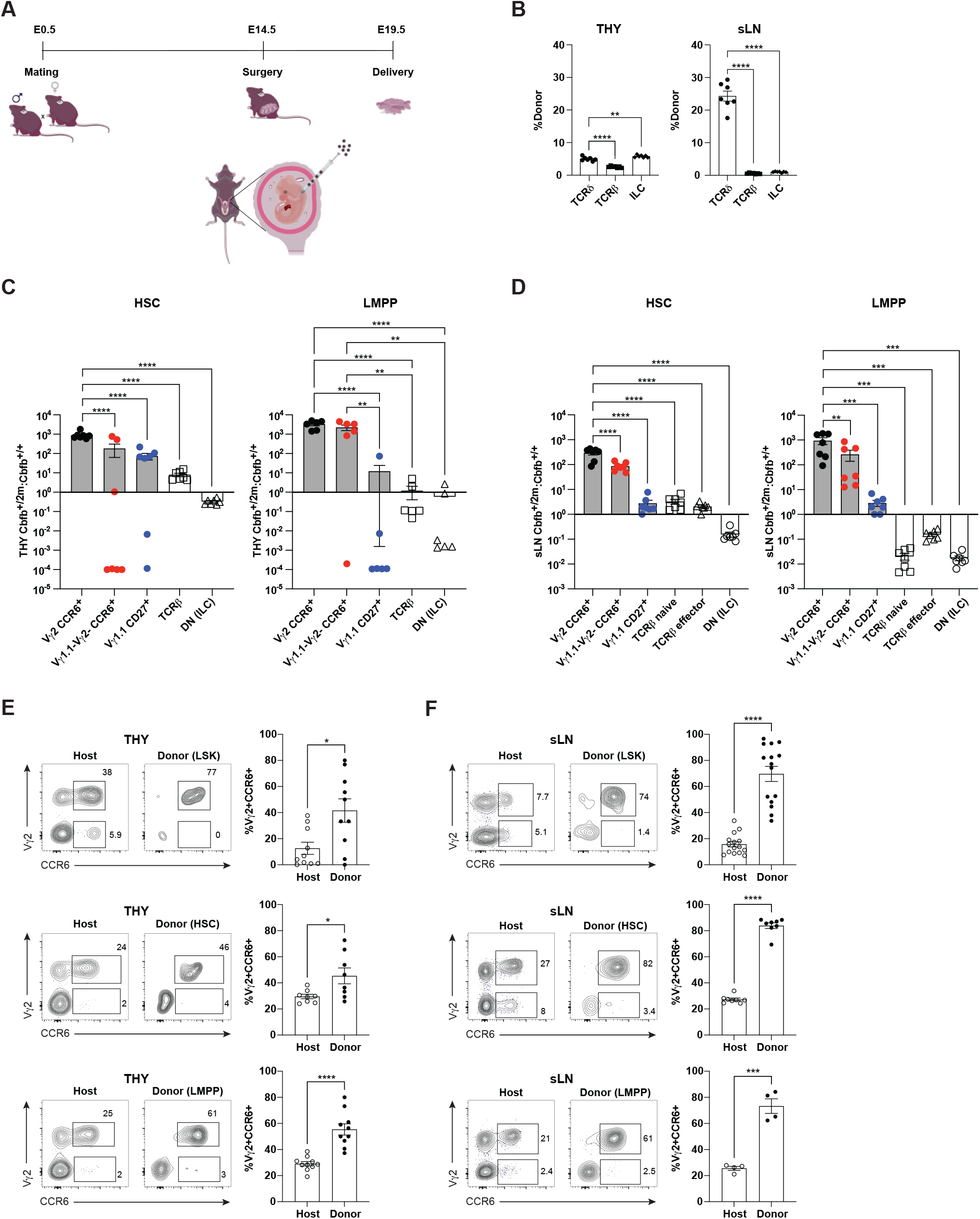
Efficient in utero differentiation of Vγ2^+^ Tγδ17 cells from adult *Cbfb*^*+/+*^ and *Cbfb*^*+/2m*^ hematopoietic progenitor subsets. (A) Experimental setup for In Utero Transplantation (IUT) assay of hematopoietic progenitor subsets in the fetal liver of E14.5 embryos. (B) Compiled data of total TCRδ^+^, TCRβ^+^ and DN (ILC) cells in neonatal THY and sLN of mice reconstituted in utero via IUT with enriched LSKs from *Cbfb*^*+/+*^ adult BM (n=7 mice per group). (C) Contribution of *Cbfb*^*+/+*^ and *Cbfb*^*+/2m*^ HSCs and LMPPs in the generation of the indicated cell subsets via mixed IUT, shown as *Cbfb*^*+/2m*^/*Cbfb*^*+/+*^ ratios, in neonatal THY (n=7 mice per group). (D) Contribution of *Cbfb*^*+/+*^ and *Cbfb*^*+/2m*^ HSCs and LMPPs in the generation of the indicated cell subsets via mixed IUT, shown as *Cbfb*^*+/2m*^/*Cbfb*^*+/+*^ ratios, in neonatal sLN (G) (n=7 mice per group). (E) Representative flow cytometric analysis and compiled data of Vγ2^+^CCR6^+^ cells in neonatal THY mice reconstituted in utero via IUT with LSKs, HSCs, or LMPPs sorted from *Cbfb*^*+/2m*^ adult BM (n?8 mice per group). (F) Representative flow cytometric analysis and compiled data of Vγ2^+^CCR6^+^ cells in neonatal sLN from mice reconstituted in utero via IUT with LSKs, HSCs, or LMPPs sorted from *Cbfb*^*+/2m*^ adult BM (n?8 mice per group). Data in (B-F) are representative of at least 3 independent experiments. Each symbol represents one mouse. *P* values determined by one-way ANOVA (B-C-D), or unpaired t-test (E-F).

We next examined whether *Cbfb2* haploinsufficient adult progenitors retain enhanced γδ T cell-generating capacity in the fetal environment compared to the WT counterparts. We generated mixed IUT chimeras transferring 50% *Cbfb*^*+/+*^ and 50% *Cbfb*^*+/2m*^ adult BM progenitors into WT hosts. The analysis of thymus in 2 weeks old pups revealed a significantly superior reconstitution of Vγ2^+^CCR6^+^ cells originating from both *Cbfb*^*+/2m*^ HSC and LMPP subsets relative to the WT counterparts in the competitive setting (**Fig. 5C**). The reconstitution of the other thymic TCRγδ^+^ IL-17–committed subset (Vγ4^+^ cells that are Vγ1.1^-^Vγ2^-^ CCR6^+^) was increased among *Cbfb*^*+/2m*^ LMPP, but not HSC-derived cells. IFNγ–producing Vγ1.1^+^ γδ T cells, TCRβ^+^ thymocytes and ILC showed variable developmental efficiency based on the progenitor subset used to reconstitute the host (**Fig. 5C**). The sLN were preferentially populated with Tγδ17 cells originating from *Cbfb*^*+/2m*^ progenitors. Conversely, WT LMPPs were the dominant source of αβ T cells and ILC in the competitive setting (**Fig. 5D**). Together, these findings demonstrate that while adult WT BM progenitors can generate Vγ2^+^ Tγδ17 cells in a fetal niche (**Fig. 5B**), reduced *Cbfb2* dosage markedly enhances this competence, conferring a selective advantage for early-life Tγδ17 cell output during physiological fetal development.

To determine whether the innate-like T cell developmental potential is confined to specific hematopoietic progenitor subsets, we transplanted sorted LSKs, HSCs, or LMPPs from adult *Cbfb*^*+/2m*^ BM into E14.5 hosts and analyzed their progenies two weeks after birth (**Fig. 5E, F**). An enhanced generation of Tγδ17 cells from all three progenitor donor subsets was observed in the thymus, where the frequencies of mature (CD24^neg^) Vγ2^+^CCR6^+^ cells, but not Vγ2^-^CCR6^+^ thymocytes (**Supp. Fig. 5E**) were higher in the donor-derived fraction (**Fig. 5E**). Remarkably, nearly all Vγ2TCR+ thymocytes from adult HSC or LMPP donors in fetal environment were Tγδ17 cells (**Fig. 5E and Supp. Fig. 5E, I)**, distinct from the generation of both CCR6^+^ and CD27^+^ Vγ2TCR+ γδ T cell subsets from HET progenitor types in adult chimeras (**Supp. Fig. 2A**). LSK progenitors were distinct, giving rise to more mature IFNγ-committed CD27^+^ Vγ2TCR+ thymocytes than CCR6^+^ Vγ2TCR+ γδ T cells (**Fig. 5C and Supp. Fig. 5I)**. These data indicate that, within the fetal environment, heterogeneous *Cbfb*^+*/2m*^ LSK progenitors contain precursors capable of generating both thymic IFNγ– and IL-17–producing Vγ2^+^ γδ T cells, whereas the more defined HSC (FLT3^-^ LSK) and LMPP (FLT3^Hi^ LSK) subsets are largely restricted to producing Vγ2^+^ Tγδ17 cells. Moreover, IFNγ–committed Vγ1.1^+^ and IL-17-committed Vγ4^+^ (Vγ1.1^-^Vγ2^-^) γδ T cells were not differently reconstituted or reduced, respectively (**Supp. Fig. 5G, K**). In sLN, the adult donor-derived cells had nearly three-fold higher frequency of Vγ2^+^CCR6^+^ γδ T cells relative to the host, and nearly two-fold higher proportion of Vγ2^+^CCR6^+^ cells compared to the host when originating from LMPP or LSK (**Fig. 5F**). In line with what we observed in the thymus, the effect of *Cbfb2* heterozygosity in sLN was muted in IL-17–committed Vγ4^+^cells and IFNγ–committed Vγ1.1^+^ cells (**Supp. Fig. 5F, H**), while haploinsufficiency drastically reduced the accumulation of both Vγ2^+^ and Vγ1.1^+^ IFNγ–committed γδ T cells (**Supp. Fig. 5J, L**). Overall, the IUT data indicate that while *Cbfb2*^+*/2m*^ adult LSK subsets uniformly favor Tγδ17 differentiation in the fetal niche, the residual potential to generate IFNγ–committed Vγ2^+^ γδ thymocytes segregates specifically with the FLT3^Int^ LSK compartment.

### *Notch1* heterozygosity in LMPPs promotes enhanced generation of Vγ2^+^ Tγδ17 cells

Unlike other γδ T cell subsets, the development of Tγδ17 cells requires NOTCH signaling via HES1^43^ and evidence supports multiple intersections between NOTCH and the RUNX:CBFβ axis^28–30^. *Notch1* expression is significantly reduced in *Cbfb*^*+/2m*^ LMPPs compared to controls (**Fig. 6A**). In contrast, other Notch receptors that are expressed at much lower levels than *Notch1* (**Supp. Fig. 6A**) are either unchanged (*Notch2* and *Notch3*) or only minimally increased (*Notch4*) in expression. Therefore, we sought to determine whether reduced NOTCH1 signaling could mimic the *Cbfb2*-dependent enhancement of Tγδ17 cell generation. To this end, we generated mice in which NOTCH1 is ablated from the LMPP stage onward by crossing *Il7ra*^*Cre*^ mice with *Notch1*^*Flox*^ mice and focused our analyses on *Il7ra*^*Cre*^:*Notch1*^*Flox/+*^ mice (LMPP^*Notch1+/-*^) mice, which lack a single copy of *Notch1* to mirror the gene-dosage reduction present in *Cbfb2* heterozygotes. Analysis of thymus and sLN from 7-day old mice revealed that LMPP^*Notch1+/-*^ mice displayed an increased generation of Vγ2^+^ Tγδ17 cells (**Fig. 6B**), accompanied by a corresponding reduction in the Vγ2^neg^CCR6^+^ cells, compared with LMPP^*Notch1+/+*^ control mice (**Supp. Fig. 6B**).

**Figure 6.**
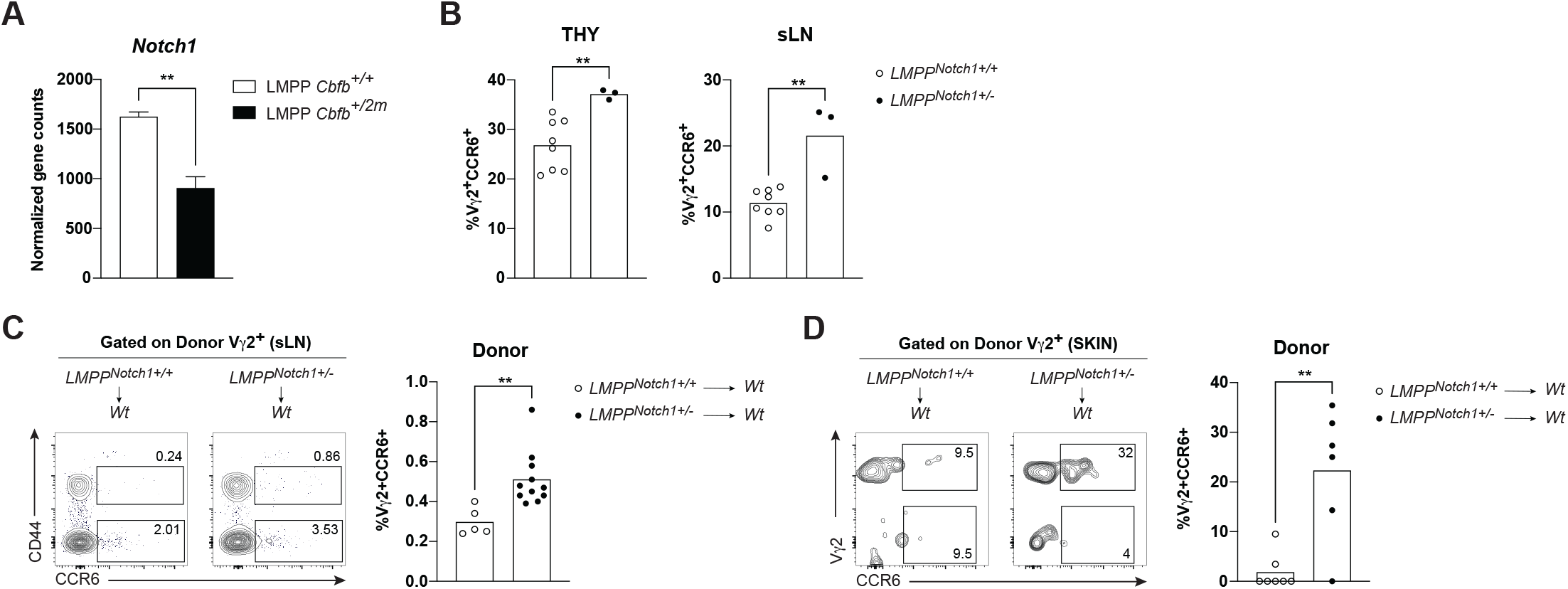
Enhanced generation of Vγ2^+^ Tγδ17 cells in *Notch1* heterozygotes. (A) *Notch1* gene expression analysis between sorted *Cbfb*^*+/+*^and *Cbfb*^*+/2m*^ BM-derived LMPPs using ULI-RNAseq gene counts. (B) Compiled frequencies of Tγδ17 cells in the thymus (top panel) and skin draining LNs (bottom panel) of 7-day old LMPP^*Notch1+/+*^ and LMPP^*Notch1+/-*^ mice (n?3 mice per group). (C) Representative flow plots and compiled frequencies of donor-derived Vγ2^+^CCR6^+^ cells from BM chimera sLN (n?5 mice per group). (D) Representative flow plots and compiled frequencies of donor-derived Vγ2^+^CCR6^+^ cells from BM chimera skin dermis (n?6 mice per group). Data in (A-C) are representative of at least 3 independent experiments. Each symbol represents one mouse. *P* values determined by unpaired t-test.

To test whether *Notch1* haploinsufficiency could endow BM progenitor with the ability to give rise to Vγ2^+^ Tγδ17 cells, we generated BM chimeras by reconstituting CD45.1 congenic hosts with adult LMPP^*Notch1+/+*^ or LMPP^*Notch1+/-*^ BM (CD45.2). In this setting, Vγ2^+^ Tγδ17 cells in sLN developed more efficiently from LMPP^*Notch1+/-*^ progenitors than from controls (**Fig. 6C**), whereas Vγ2^neg^CCR6^+^ cells were unaffected (**Supp. Fig. 6C**). We also observed increased frequencies of Vγ2^+^ Tγδ17 cells derived from LMPP^*Notch1+/-*^ BM in the skin of chimeric mice compared to controls (**Fig. 6D**). Consistent with the sLN analysis, Vγ2^neg^CCR6+ cells in the skin were unaffected (**Supp. Fig. 6C**). All together, these findings point to a *Notch1* dose-dependent mechanism that promotes the generation of Vγ2^+^ Tγδ17 cells, resembling *Cbfb2* heterozygosity, and supporting a direct quantitative regulation of *Notch1* by CBFβ2 complex.

## Discussion

Our study identifies *Cbfb2* gene dosage as a central quantitative regulator of fetal versus adult lymphoid lineage potential in the T cell lineage. We demonstrate that reduction in CBFβ2 levels is sufficient to reveal fetal-like lymphoid developmental competence in adult hematopoietic progenitors differentiating in an adult environment. The adult BM progenitors generated Vγ2^+^ Tγδ17 cells fully functional as they populated the skin and responded robustly in models of psoriasis-like dermatitis and *Malassezia furfur*-induced skin inflammation. These findings challenge the prevailing view that the loss of innate-like T cell lineage potential in adulthood is irreversible and that differences in fetal and adult lymphopoiesis are established by distinct waves of progenitors with fundamentally different gene circuit hardwiring. Rather, our results support a model in which fetal versus adult developmental programs are maintained through quantitative tuning of core transcriptional circuits in progenitors whose developmental potentials are modulated by environmental factors.

A key advance of this work is the demonstration that adult LMPPs carrying a 50% reduction in CBFβ2, without disruption of RUNX proteins themselves, acquire the capacity to generate functional Vγ2^+^ Tγδ17 cells that represent prototypical innate-like T cells, both in adult BM chimeras and within a physiological fetal niche via IUT. Moreover, adult WT progenitors show capacity to generate Tγδ17 cells in the fetal environment, albeit much less efficiently than adult *Cbfb*^*+/2m*^ progenitors, demonstrating robust developmental plasticity and emphasizing the importance of developmental niche in progenitor differentiation trajectory. Whether the “fetalness” can be primarily instituted by dynamic modulation of the RUNX:CBFβ transcriptional module in adult progenitors by fetal factors remains to be determined.

Mechanistically, the impact of *Cbfb2* haploinsufficiency cannot be explained solely by steady-state transcriptional changes. Adult *Cbfb*^*+/2m*^ HSCs, LMPPs, and CLPs displayed only modest transcriptional alterations, with LMPPs showing the clearest but still limited differences. Instead, chromatin profiling revealed that reduced CBFβ2 dosage reconfigures the epigenetic landscape of adult LMPPs toward a fetal-like state, including broad gains of H3K4me3 at loci associated with lymphopoiesis and hematopoietic development. These findings support a model in which CBFβ2 quantity regulates chromatin accessibility and lineage priming, enabling adult progenitors to access developmental trajectories normally restricted to fetal hematopoiesis. The amplification of Tγδ17 cell generation in IUT experiments supports a two-step model: CBFβ2 dosage establishes progenitor competence through epigenetic priming; and the fetal microenvironment provides permissive cues, possibly via NOTCH ligands, cytokines, or stromal interactions, that actualize this potential. Such synergy explains how subtle quantitative differences in a ubiquitous transcriptional cofactor can produce stage-specific developmental outcome. Further studies will be required to definitively test this hypothesis.

We further identify convergence between CBFβ2-mediated regulation and NOTCH1 signaling in innate-like T cell development. Cooperation of RUNX1 and NOTCH signaling in T cell lineage commitment is well established^45^. While genetic studies have indicated NOTCH1 regulation of *Cbfb* expression^28,46,47^, data also suggest a bidirectional regulation during T cell development and transformation whereby RUNX:CBFβ complex in turn modulates *Notch1* expression^48,49^. *Notch1* heterozygosity phenocopied *Cbfb2* haploinsufficiency, enhancing Vγ2^+^ Tγδ17 generation and barrier-tissue colonization. Given that NOTCH1 can regulate *Cbfb* transcription and shares downstream targets with RUNX:CBFβ complexes, reduced NOTCH1 signaling may lower functional *Cbfb2* output or relax RUNX-dependent lineage restriction. Recent work further demonstrated that NOTCH signaling can drive a functional conversion of RUNX transcription factors, promoting protein complex reorganization and genomic redeployment, to initiate the T-lineage program^30^. This mechanistic insight provides a compelling explanation for how quantitative reductions in NOTCH1 could reshape RUNX-dependent chromatin landscapes, thereby aligning with the fetal-like lineage trajectories observed in both *Cbfb2* and *Notch1* haploinsufficient progenitors. The parallel outcomes of *Cbfb2* and *Notch1* haploinsufficiency supports a bidirectional regulatory mechanism whereby quantitative tuning of core transcriptional circuits, rather than binary on/off switches, underlies age-dependent lymphoid potential.

## Material and Methods

### Mice

C57BL/6J mice (Stock no: 000664), CD45.1 mice (Stock no: 002014), *Thy1*.*1* mice (Stock no: 000406), and *Notch1*^*Flox*^ mice (Stock no: 006951) were from Jax Laboratories. *Cbfb*^*2m*^ mice and *Il7ra*^*Cre*^ mice were previously described^24,50^. Males and females were used for experiments, but sex-matched withinan experiment. No differences were observed between sexes. Ages of mice used for experiments are indicated in Figure Legends. Animals were randomly allocated to experimental groups. All mouse procedures were approved by the University of Massachusetts Chan Medical School IACUC.

### Sample preparation and flow cytometry

Skin draining (cervical, inguinal, axillary and brachial) lymph nodes (sLN), thymus (THY), and fetal liver (FL) were collected into complete media (RPMI, 2% FCS, 10 mM HEPES, 1% Pen/Strep). The tissues were then smashed through a 70 µm cell strainer. Bone marrow (BM) cells were harvested by flushing both tibia and femurs and filtered through a 70 µm cell strainer. Isolated cells were then washed with FACS buffer (DPBS, 2% FCS, 2 mM EDTA). Skin single cell suspensions were prepared from mouse ears. As previously described^39^, ears were depilated using Nair application for two minutes, followed by gentle removal of cream and rinsing of tissues using PBS. Depilated mouse ears were then peeled into dorsal and ventral halves, chopped finely using scissors, and digested with1 U/mL Liberase TL (Roche) + 0.5 mg/mL Hylauronidase (Sigma-Aldrich) + 0.05 mg/mL DNAse (Roche) dissolved in skin digestion buffer (HBSS, 5% FCS, 1 mM HEPES) on a stir plate for 90 minutes at 37°C. To stop digestion, 10 mM EDTA (Teknova) was added. Cells were filtered through 100 µm cell strainer, rinsed with skin digestion buffer, spun down, filtered through 70 µm cell strainer, rinsed again, spun down, and resuspended. Flow cytometry staining was performed in 96-well microtiter round-bottom plates. Antibody cocktails were diluted in FACS buffer and cells were stained in 50 µL for 20 min on ice. All the antibodies were purchased from BD Biosciences, BioLegend or eBioscience: CD3ε (500A2), CD4 (RM4-5), CD8α (53-6.7), CD8β (YTS156.7.7), TCRβ (H57-597), TCRδ (GL3), Vγ2 (UC3-10A6), Vγ1.1 (2.11), CD27 (LG.3A10), CCR6 (140706), CD24 (M1/69), CD44 (IM7), CD25 (PC61), CD45 (30-F11), IL-17A (7B7), NK1.1 (PK136), Gr1 (RB6-8CS), CD11b (M1/70), CD11c (N418), TER119 (TER-119), CD19 (ID3), CD45.1 (A20), CD45.2 (104), THY1.1 (OX-7), CD34 (RAM34), c-Kit (2B8), Sca1 (D7), IL-7Rα (A7R34), FLT3 (A2F10), streptavidin PerCp-Cyanine5.5, and streptavidin BV421. 5-OP-RU and 6-FP–loaded mMR1 tetramers were provided by the NIH Tetramer Core Facility. Cell suspensions were stained with FixableViability Dye eFluor780 or eFluor510 (eBioscience) in FATHCS buffer to exclude dead cells from all analysis, and Fc receptors blocked with anti-mouse CD16/32. For intracellular cytokine staining, cells were either stimulated in vitro with 10 ng/mL phorbol myristate acetate (PMA) + 1 mM Ionomycin (both Sigma-Aldrich), in the presence of GolgiPlug and GolgiStop (both BD Biosciences) for 4 hours at 37°C or incubated with GolgiPlug and GolgiStop for 4 hours at 37°C. Cells were then surface stained, fixed/permeabilized with Cytofix/Cytoperm buffer (BD Biosciences) and then stained for indicated intracellular cytokines. Cells were incubated for 20 min on ice with antibodies for surface staining, 1 hour at room temperature for tetramer staining, and 30 min on ice for intracellular staining. Data were collected on a BD LSR II or FACSymphony A5 and analyzed in FlowJo v10 software.

### Cell sorting

Hematopoietic progenitor cells were isolated from either E14.5 FL or adult BM. Cells were stained as described above and sorted on a BD Fusion with an 85 µm nozzle. LSKs were sorted as live Lin^-^ c-Kit^+^ Sca1^+^, HSCs as live Lin^-^ c-Kit^+^ Sca1^+^ FLT3^-^, LMPPs as live Lin^-^ c-Kit^+^ Sca1^+^ FLT3^High^, and CLPs as live Lin^-^ c-Kit^Int^ Sca1^Int^ FLT3^+^ IL-7Ra^+^. Cells were maintained at 4°C until ready for sorting.

### Bone marrow chimeras, retroviral transduction

CD45.1 mice were lethally irradiated twice with 550 rads gamma-irradiation, 3 hours apart, then intravenously injected with 1-3 x 10^6^ bone marrow cells. Mice were analyzed 6-8 weeks after cell transfer. For retroviral transduction, PlatE cells were transfected with murine stem cell virus retroviral constructs encoding full length *Cbfb2* with Lipofectamine 2000 following manufacturer’s protocol. For transduction of BM-derived cells, BM was harvested 4 days after 5-flurouracil injection and cultured in the presence of recombinant IL-3 (20 µg/ml), IL-6 (50 µg/ml), and mouse stem cell factor (100 ng/ml). BM cells were spin-infected twice with a retroviral construct expressing *Cbfb2* with THY1.1 as a reporter. One day after the last spin infection, the cells were injected into lethally irradiated CD45.1 recipients. Mice were analyzed 6-8 weeks later.

### In Utero Transplantation (IUT) assay

Embryonic day (E)14.5 CD45.1 fetuses were injected with FACS-sorted LSKs, HSCs, or LMPPs. Donor cells were equally distributed among all injected fetuses. The injections were performed directly into the fetal liver (5 µl per fetus) using in-house pulled glass micropipettes and a microinjector, as previously described^41^. Briefly, pregnant dams were anesthetized, and a laparotomy was performed to expose the uterus. Individual fetuses were injected through the intact, translucent uterine wall. The laparotomy was closed in layers, and dams were allowed to recover and deliver at term few days later. Pups were analyzed 2-3 weeks after birth.

### Imiquimod-induced psoriasis-like dermatitis

6 weeks after reconstitution, BM chimeric mice were anesthetized with isoflurane and treated topically on each ear with 5 mg of 5% imiquimod cream (Imiquimod Cream 5%; Perrigo) or petroleum jelly (vehicle) once daily for up to 5 consecutive days. Peripheral and central ear thicknesses were measured daily with a digital caliper (Mitutoyo). Vehicle-treated and M. furfur–associated mice were housed separately to prevent cross-contamination.

### Epicutaneous association of mice with *Malassezia furfur*

*M. furfur* strain (Robin) Baillon (CBS 14521) was grown in modified Dixon (mDixon) medium at 30°C and 180 rpm for 2-3 days. Cells were washed in PBS and suspended in commercially available native olive oil at a density of 2 OD_A600_/ml. A 100 μL suspension (corresponding to 2 OD_A600_ ≈ 1 x 10^6^ yeast cells) was topically applied once to the dorsal ear skin of anesthetized BM chimeric mice, 6 weeks post-reconstitution. Control animals received olive oil (vehicle) alone. Vehicle-treated and *M. furfur*-associated mice were housed separately to prevent cross-contamination. Mice were analyzed 7 days after association.

### UltraLowInput (ULI)-RNAseq and computational analysis

After the final sort of 1,000 cells directly into 5 µl lysis buffer (TCL buffer with 1% 2-Mercaptoethanol), Smart-seq2 libraries were prepared as previously described^51,52^ with slight modifications and according to ImmGen.org Standard Operating Procedures. Briefly, total RNA was captured and purified on RNAClean XP beads (Beckman Coulter). Polyadenylated mRNA was then selected using an anchored oligo(dT) primer (5’-AAGCAGTGGTATCAACGCAGAGTACT30VN-3’) and converted to cDNA via reverse transcription. First-strand cDNA was subjected to limited PCR amplification followed by Tn5 transposon-based fragmentation using the Nextera XT DNA Library Preparation Kit (Illumina). Samples were then PCR-amplified for 18 cycles using barcoded primers such that each sample carries a specific combination of eight base Illumina P5 and P7 barcodes and pooled together prior to sequencing. Paired-end sequencing was performed on an Illumina NextSeq500 using 2 x 25bp reads.

Reads were aligned to the mouse genome (GENCODE GRCm38/mm10 primary assembly and gene annotations vM25; https://www.gencodegenes.org/mouse/release_M25.html) with STAR 2.7.3a^53^. The ribosomal RNA gene annotations were removed from GTF (General Transfer Format) file. The gene-level quantification was calculated by featureCounts. Raw reads count tables were normalized by median of ratios method with DESeq2 package from Bioconductor^54^ and then converted to GCT and CLS format.

Samples with less than 1 million uniquely mapped reads, or having less than 8,000 genes with >10 reads, or with Transcript Integrity score <45 were removed from the data set prior to downstream analysis and excluded from normalization to mitigate the effect of poor-quality samples on normalized counts. All samples were also screened for contamination by using known cell-type-specific transcripts (per ImmGen ULI-RNAseq and microarray data). In practice, the acceptable threshold was set at 0.1 of typical gene expression of contaminant cell types. Retaining such samples can create structure in the data, and/or generate false distances between samples. In addition, biological replicates were analyzed for Pearson correlation to identify poor-quality samples and remove them from the data set. Pearson correlation was calculated on transcripts with an average of >5 reads or below the 99th percentile for number of reads in the dataset to avoid outlier effects. Any replicates that did not exhibit a correlation of 0.9 or greater were removed from the data set prior to downstream analysis. Finally, the RNA integrity for all samples was measured by median Transcript Integrity across mouse housekeeping genes with RSeQC 2.6.4^55^.

Differential gene expression analysis was performed using normalized gene expression values. Statistical significance was assessed using DESeq2, and resulting P values were adjusted for multiple testing. Principal component analysis (PCA) was performed on normalized gene expression data to visualize global transcriptional differences between samples.

## CUT&RUN

A low-input (10,000 primary cells) of the CUT&RUN technique^56^ developed by ImmGen.org in close collaboration with Epicypher was applied here. Briefly, hematopoietic progenitor subsets were prepared and sorted as above, except that a LIVE/DEAD Fixable Aqua Dead Cell Stain (Thermo Fisher Scientific) was used as a viability marker. After staining, cell suspensions were washed and lightly fixed in 200 μl of 0.1% formaldehyde in PBS (freshly diluted from 37% stock; Sigma-Aldrich) for exactly 1 min at room temperature, before quenching with 10 μl 2.5 M glycine in 200 μl. Cells were then washed in FACS medium, and 8 × 104 Bfo cells were sorted, with the same gating strategies as above. Sort volume was measured and diluted with an equal volume of 2X Nuclei Extraction buffer [40 mM Hepes, 20 mM KCl, 0.2% Triton X-100, 40% Glycerol, 2 mM DTT, 1 mM Spermidine, 2X Roche Complete Protease Inhibitor (Millipore Sigma)] and freshly supplemented to 2X KDAC inhibitor cocktail (2 μM trichostatin A, 1 mM sodium butyrate, 1 mM nicotinamide in 70% DMSO). Samples were then frozen in an isopropanol-filled container at -80°C. Samples were thawed on ice and diluted to 10^5^ cells/ml in 1X Nuclei Extraction buffer. 10 μl of activated Concanavalin A beads, 2 μl of 1:50 (vol/vol) SNAP-CUTANA K-MetStat Panel, and 0.5 µg of primary antibody [rabbit IgG (EpiCypher), H3K4me3 (EpiCypher)] were added per reaction (10^4^ cells) and incubated overnight. The next day, the beads were washed in 250 μl Digitonin Buffer (20 mM pH 7.5 Hepes, 150 mM NaCl, 0.5 mM Spermidine, 1X Roche Complete mini, 0.01% digitonin) twice prior to the addition of 5 μl CUTANA pAG-MNase (20X) in 50 μl of Digitonin Buffer per reaction. Beads were again washed twice in 250 μl Digitonin Buffer and suspended in 50 μl Digitonin Buffer. For chromatin digestion, CaCl_2_was added to 2 mM in each reaction to activate the MNase. 33 μl of High Salt Stop Buffer (750 mM NaCl, 26.4 mM EDTA, 5.28 mM EGTA, 66 µg/ml RNase A, 66 µg/ml Glycogen) was added to each reaction to terminate the MNase activity after a 2-h incubation at 4°C. 20 pg CUTANA *E. coli* Spike-in DNA (EpiCypher) was added per sample. Samples were incubated for 10 mins at 37°C to release the cleaved chromatin. CUT&RUN-enriched DNA was isolated from the Concanavalin A beads and cleaned up using 2:1 (Bead:DNA) ratio of Serapure beads. Libraries were prepared using a CUTANA CUT&RUN Library Prep Kit (EpiCypher) and sequenced on Illumina NextSeq 1000/2000 (paired end 2 x 100 bp read). Fastq files were QC and trimmed using FASTQC and Trimmomatic. Samples were aligned to mm10 reference genome using Bowtie2, sorted and filtered with SAMTools. Normalization was performed using sequencing depth of each sample and spiked-in *E. coli* seq depth. Peak calling was performed using SEACR^57^ to adjust for narrow peaks (H3K4me3). Differential peaks analysis was performed using DiffBind package to generate affinity binding matrix. Replicates of each sample were used for differential analysis with DESeq2. Significantly differential peaks were defined by P <0.05.

### Quantification and statistical analysis

Statistical treatment of the gene expression data is as described above. Summary data from FACS analyses were analyzed in GraphPad Prism software using statistical tests indicated in Figure Legends. The mean of all samples in a group is used to represent the central tendency of the dataset, and all error bars represent SEM of biological replicates. Sample size was not determined prior to experimentation. The number of biological replicates (n) is stated in the Figure Legends, and P < 0.05 was considered significant. No randomization of experiments was conducted. Experimenters were not blinded during performance or analysis of the experiments.

