## Supplemental File for "*Cbfb2* gene dosage programs the differential lymphoid lineage developmental potential of fetal and adult hematopoietic progenitors"

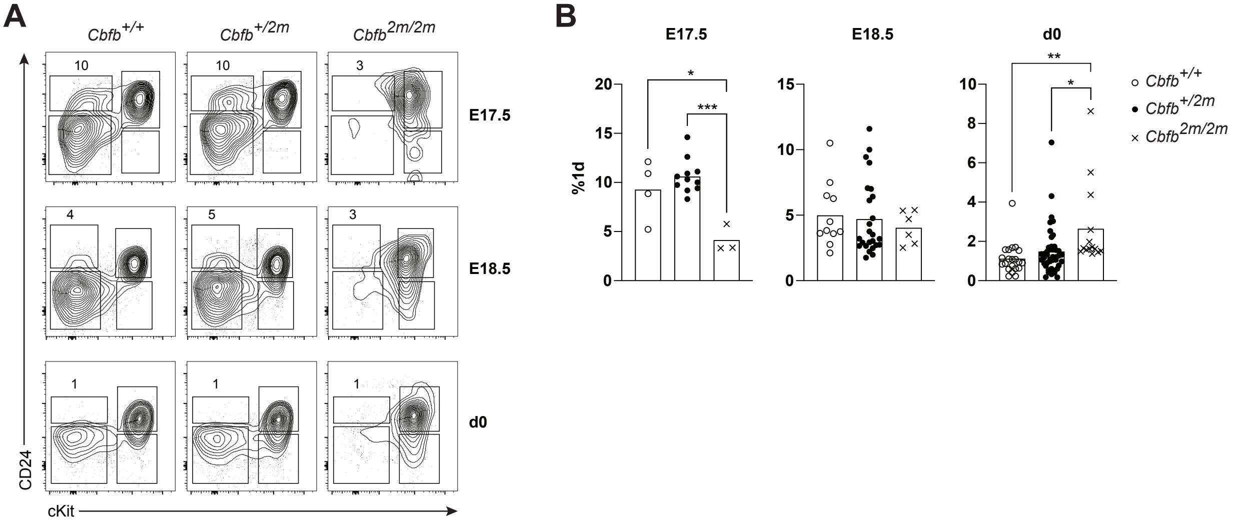

A
Cbfb+/+
Cbfb+/2m
Cbfb2m/2m
10
10
3
E17.5
4
5
3
E18.5
1
1
1
d0
CD24
cKit
B
E17.5
E18.5
d0
%1d
*
***
**
*
Cbfb+/+
Cbfb+/2m
Cbfb2m/2m

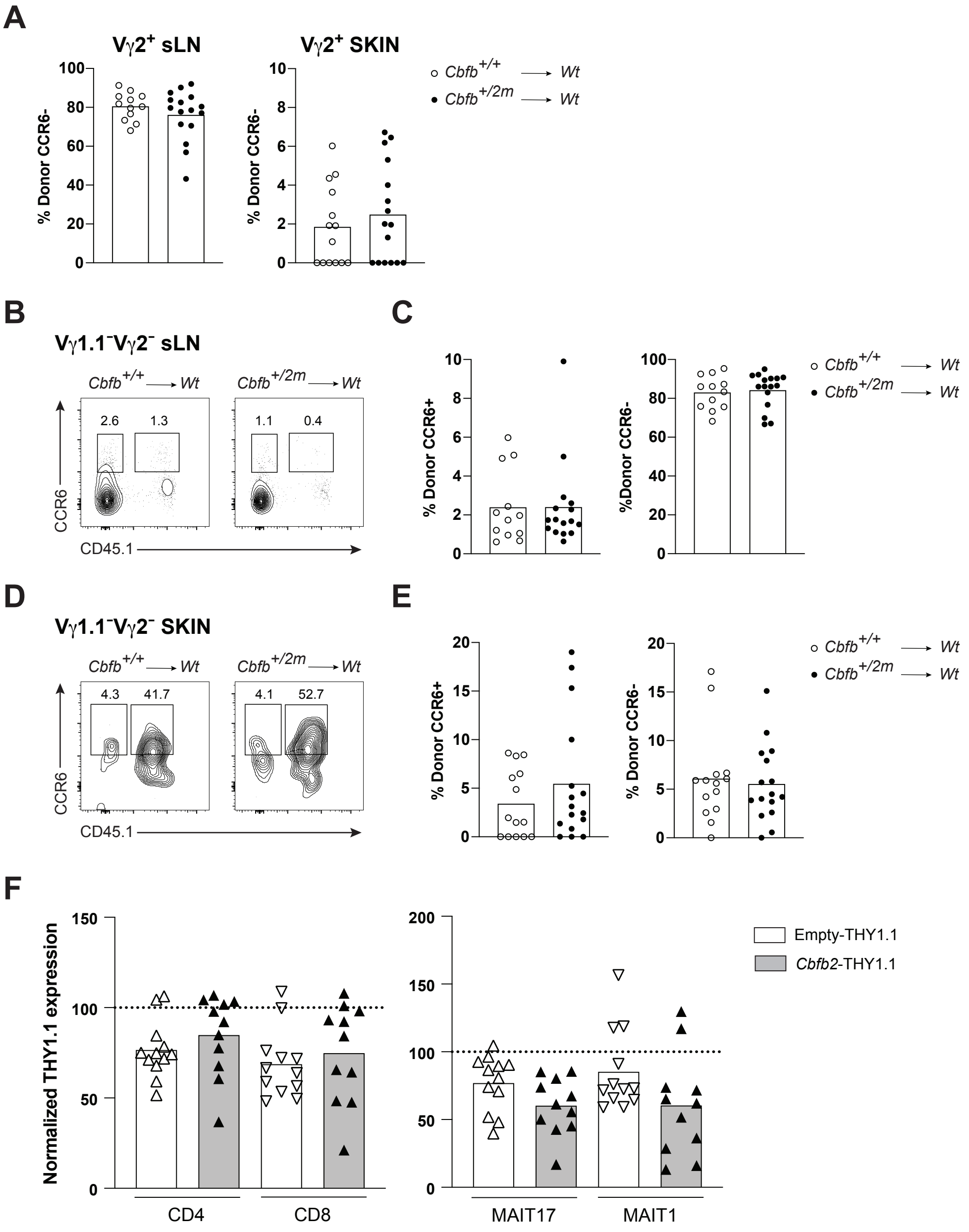

A
Vγ2+ sLN
Vγ2+ SKIN
% Donor CCR6-
Cbfb+/+ → Wt
Cbfb+/2m → Wt
B
Vγ1.1−Vγ2− sLN
Cbfb+/+ → Wt
Cbfb+/2m → Wt
2.6
1.3
1.1
0.4
CCR6
CD45.1
C
% Donor CCR6+
%Donor CCR6-
D
Vγ1.1−Vγ2− SKIN
4.3
41.7
4.1
52.7
E
% Donor CCR6+
% Donor CCR6-
F
Normalized THY1.1 expression
Empty-THY1.1
Cbfb2-THY1.1
CD4
CD8
MAIT17
MAIT1

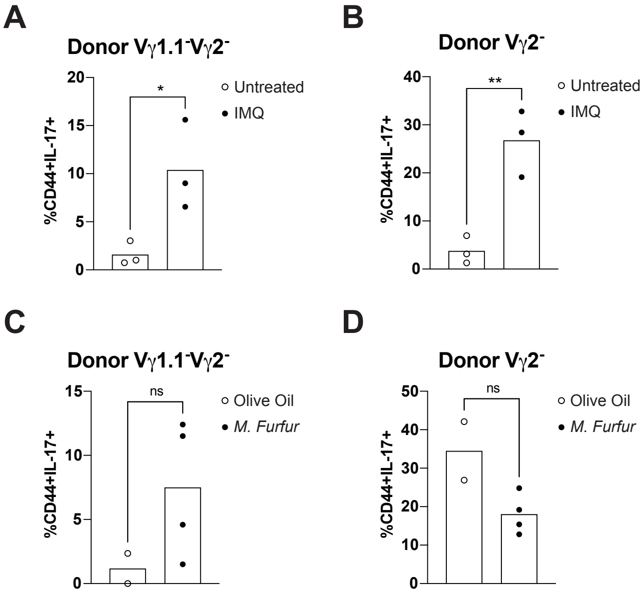
A
Donor Vγ1.1⁻Vγ2⁻
%CD44+IL-17+
Untreated
IMQ
*
B
Donor Vγ2⁻
%CD44+IL-17+
Untreated
IMQ
**
C
Donor Vγ1.1⁻Vγ2⁻
%CD44+IL-17+
Olive Oil
M. Furfur
ns
D
Donor Vγ2⁻
%CD44+IL-17+
Olive Oil
M. Furfur
ns

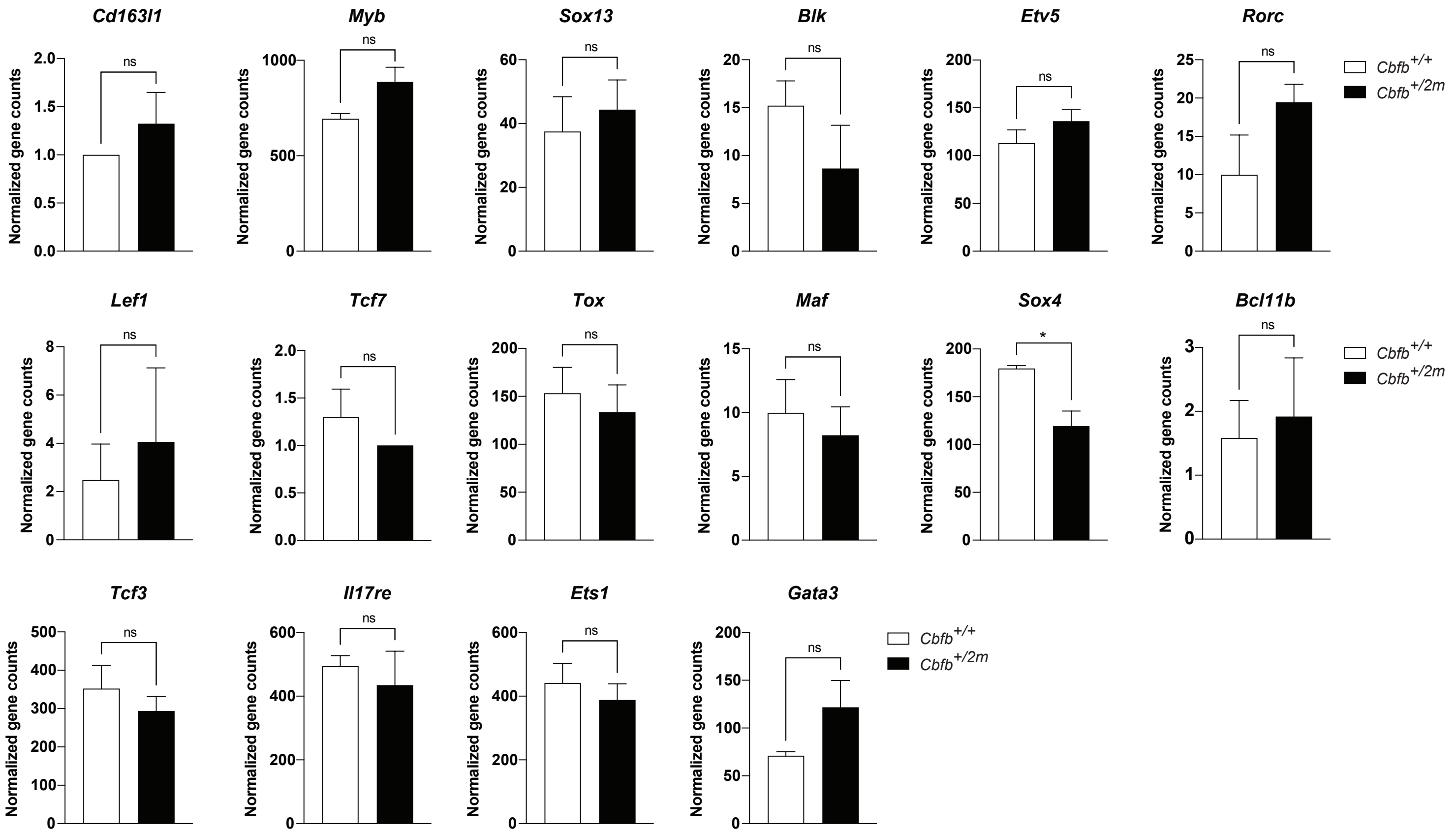

Cd163l1
Myb
Sox13
Blk
Etv5
Rorc
Lef1
Tcf7
Tox
Maf
Sox4
Bcl11b
Tcf3
Il17re
Ets1
Gata3
Normalized gene counts
ns
*
Cbfb+/+
Cbfb+/2m

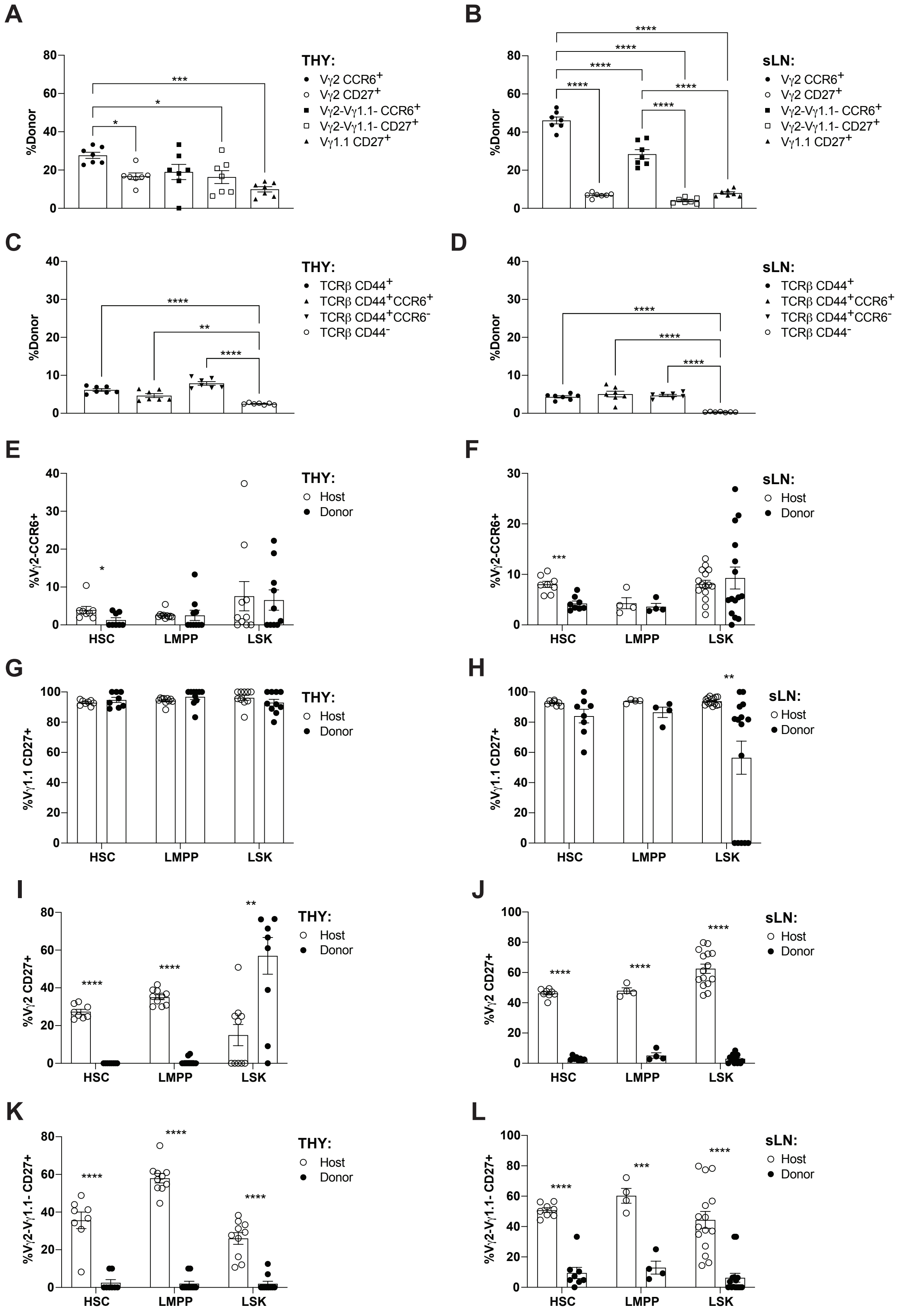

A
B
C
D
E
F
G
H
I
J
K
L
%Donor
THY:
sLN:
Vγ2 CCR6+
Vγ2 CD27+
Vγ2-Vγ1.1- CCR6+
Vγ2-Vγ1.1- CD27+
Vγ1.1 CD27+
TCRβ CD44+
TCRβ CD44+CCR6+
TCRβ CD44+CCR6-
TCRβ CD44-
Host
Donor
%Vγ2-CCR6+
%Vγ1.1 CD27+
%Vγ2 CD27+
%Vγ2-Vγ1.1- CD27+
HSC
LMPP
LSK

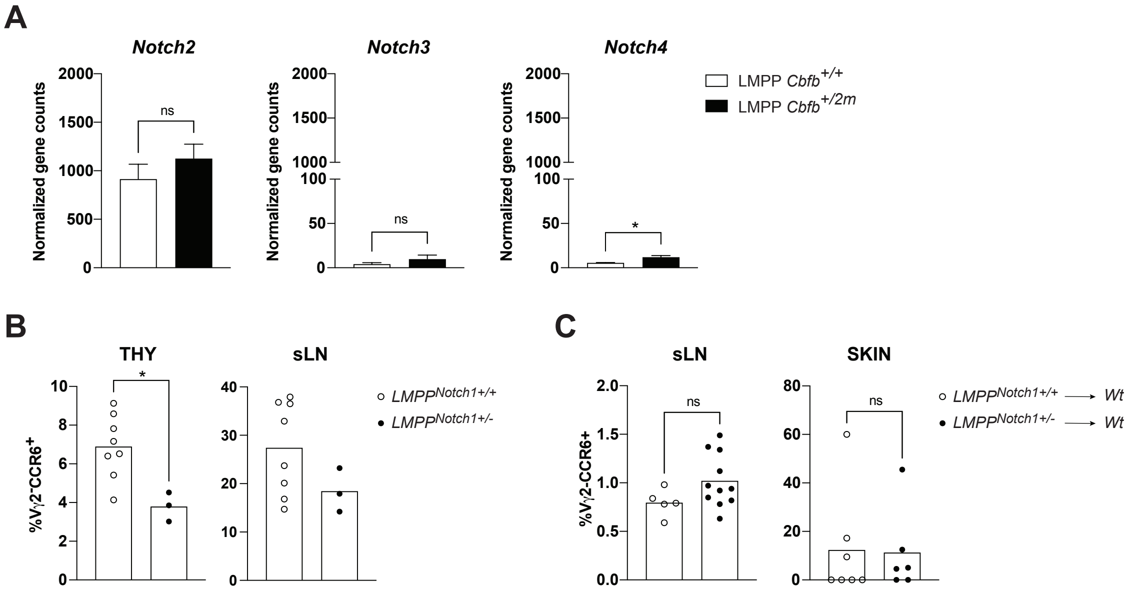
A
Notch2
Notch3
Notch4
Normalized gene counts
2000
1500
1000
500
0
100
50
ns
*
LMPP Cbfb+/+
LMPP Cbfb+/2m
B
THY
sLN
%Vγ2-CCR6+
LMPPNotch1+/+
LMPPNotch1+/-
C
sLN
SKIN
%Vγ2-CCR6+
LMPPNotch1+/+ → Wt
LMPPNotch1+/- → Wt

**Supplementary Figure Legends**

**Figure S1. Dose-dependent reduction of DN1d thymic precursors in *Cbfb2* mutant mice during late fetal development and birth. Related to Figure 1.**

(A) Representative flow cytometric contour plots depicting expression of CD24 and cKit in DN1 ($CD44^+CD25^-$ DN) thymocytes from $Cbfb^{+/+}$, $Cbfb^{+/2m}$, and $Cbfb^{2m/2m}$ mice analyzed at E17.5, E18.5 and day 0 (delivery day).

(B) Summary of the compiled flow cytometric data of A (n≥3 mice per genotype).

Data in (A-B) are representative of at least 3 independent experiments. Each symbol represents one mouse. *P* values determined by two-way ANOVA.

**Figure S2. *Cbfb*$^{+/2m}$ bone marrow fails to enable Vγ1.1$^-$Vγ2$^-$ Tγδ17 development, and ectopic expression of CBFβ2 does not perturb αβT or MAIT cell output. Related to Figure 2.**

(A) Compiled frequencies of $CCR6^-$ donor cells among Vγ2$^+$ cells in BM chimera sLN and skin dermis (n≥12 mice per group).

(B) Representative flow plots and compiled frequencies for Vγ1.1$^-$Vγ2$^-$ cells in BM chimera sLN.

(C) Compiled frequencies of $CCR6^+$ and $CCR6^-$ donor cells among Vγ1.1$^-$Vγ2$^-$ cells in BM chimera from (B) (n≥12 mice per group).

(D) Representative flow plots and compiled frequencies for Vγ1.1$^-$Vγ2$^-$ cells in BM chimera skin dermis.

(E) Compiled frequencies of $CCR6^+$ and $CCR6^-$ donor cells among Vγ1.1$^-$Vγ2$^-$ cells in BM chimera from (D) (n≥12 mice per group).

(F) Summary of αβT conventional and MAIT subsets after normalization of THY1.1 expression (n≥11 mice per group).

Data in (A-F) are representative of at least 3 independent experiments. Each symbol represents one mouse. *P* values determined by unpaired t-test.

**Figure S3. Effector function analysis of Vγ1.1$^-$Vγ2$^-$ Tγδ17 derived from *Cbfb*$^{+/2m}$ BM. Related to Figure 3.**

(A) Compiled frequencies of IL-17 secretion among donor Vγ1.1$^-$Vγ2$^-$ cells in BM chimera sLN after IMQ treatment (n=3 mice per group).

(B) Compiled frequencies of IL-17 secretion among donor Vγ2$^-$ cells in BM chimera skin dermis after IMQ treatment (n=3 mice per group).

(C) Compiled frequencies of IL-17 secretion among donor Vγ1.1$^-$Vγ2$^-$ cells in BM chimera sLN after fungal colonization (n≥2 mice per group).

(D) Compiled frequencies of IL-17 secretion among donor Vγ2$^-$ cells in BM chimera skin dermis after fungal colonization (n≥2 mice per group).

Each symbol represents one mouse. *P* values determined by unpaired t-test.

**Figure S4. Analysis of the Vγ2$^+$ Tγδ17 gene signature. Related to Figure 4.**

Differential gene expression analysis between sorted $Cbfb^{+/+}$and $Cbfb^{+/2m}$ BM-derived LMPPs using ULI-RNAseq gene counts.

Samples were analyzed in biological triplicates. *P* values determined by unpaired t-test.

**Figure S5. In utero differentiation potential of adult *Cbfb*$^{+/2m}$ hematopoietic progenitor subsets. Related to Figure 5.**

(A) Compiled data of TCRδ$^+$ cell subsets in neonatal THY of mice reconstituted in utero via IUT with enriched LSKs from $Cbfb^{+/+}$ adult BM (n=7 mice per group).

(B) Compiled data of TCRδ$^+$ cell subsets in neonatal sLN of mice reconstituted in utero via IUT with enriched LSKs from $Cbfb^{+/+}$ adult BM (n=7 mice per group).

(C) Compiled data of TCRβ$^{+}$ cell subsets in neonatal THY of mice reconstituted in utero via IUT with enriched LSKs from *Cbfb*$^{+/+}$ adult BM (n=7 mice per group).
(D) Compiled data of TCRβ$^{+}$ cell subsets in neonatal sLN of mice reconstituted in utero via IUT with enriched LSKs from *Cbfb*$^{+/+}$ adult BM (n=7 mice per group).
(E) Compiled data of Vγ2$^{-}$CCR6$^{+}$ among TCRd$^{+}$ cells in neonatal THY from mice reconstituted in utero via IUT with sorted LSKs, HSCs, or LMPPs from *Cbfb*$^{+/2m}$ adult BM (n≥8 mice per group).
(F) Compiled data of Vγ2$^{-}$CCR6$^{+}$ among TCRd$^{+}$ cells in neonatal sLN from mice reconstituted in utero via IUT with sorted LSKs, HSCs, or LMPPs from *Cbfb*$^{+/2m}$ adult BM (n≥4 mice per group).
(G) Compiled data of CD27$^{+}$ cells among Vγ1.1$^{+}$ in neonatal THY from mice reconstituted in utero via IUT with sorted LSKs, HSCs, or LMPPs from *Cbfb*$^{+/2m}$ adult BM (n≥8 mice per group).
(H) Compiled data of CD27$^{+}$ cells among Vγ1.1$^{+}$ in neonatal sLN from mice reconstituted in utero via IUT with sorted LSKs, HSCs, or LMPPs from *Cbfb*$^{+/2m}$ adult BM (n≥4 mice per group).
(I) Compiled data of CD27$^{+}$ cells among Vγ2$^{+}$ in neonatal THY from mice reconstituted in utero via IUT with sorted LSKs, HSCs, or LMPPs from *Cbfb*$^{+/2m}$ adult BM (n≥8 mice per group).
(J) Compiled data of CD27$^{+}$ cells among Vγ2$^{+}$ in neonatal THY from mice reconstituted in utero via IUT with sorted LSKs, HSCs, or LMPPs from *Cbfb*$^{+/2m}$ adult BM (n≥4 mice per group).
(K) Compiled data of CD27$^{+}$ cells among Vγ1.1$^{-}$Vγ2$^{-}$ in neonatal THY from mice reconstituted in utero via IUT with sorted LSKs, HSCs, or LMPPs from *Cbfb*$^{+/2m}$ adult BM (n≥8 mice per group).
(L) Compiled data of CD27$^{+}$ cells among Vγ1.1$^{-}$Vγ2$^{-}$ in neonatal sLN from mice reconstituted in utero via IUT with sorted LSKs, HSCs, or LMPPs from *Cbfb*$^{+/2m}$ adult BM (n≥4 mice per group).

Data in (A-H) are representative of at least 3 independent experiments. Each symbol represents one mouse. *P* values determined by unpaired t-test.

**Figure S6. *Notch1* haploinsufficiency selectively alters LMPP Notch signaling without promoting Vγ2$^{-}$CCR6$^{+}$ Tγδ17 cells. Related to Figure 6**.

(A) Differential gene expression analysis between sorted *Cbfb*$^{+/+}$and *Cbfb*$^{+/2m}$ BM-derived LMPPs using ULI-RNAseq gene counts.
(B) Compiled frequencies of Vγ2$^{-}$CCR6$^{+}$ cells in the thymus and skin draining LNs of 7-day old LMPP$^{Notch1+/+}$ and LMPP$^{Notch1+/-}$ mice (n≥3 mice per group).
(C) Compiled frequencies of donor-derived Vγ2$^{-}$CCR6$^{+}$ from BM chimera sLN and skin dermis (n≥5 mice per group).

Data in (A-C) are representative of at least 3 independent experiments. Each symbol represents one mouse. *P* values determined by unpaired t-test.
